# Sex differences in GLP-1 signaling across species

**DOI:** 10.1101/2025.03.17.643822

**Authors:** Thomas Roseberry, Irene Grossrubatscher, Tim Krausz, Yuliang Wang, Michael Schwartz, David Tingley

## Abstract

Two billion humans are currently overweight or obese^1^. While glucagon-like peptide 1 receptor (GLP1R) agonists have emerged as the most promising treatment for this epidemic, side effects including nausea and vomiting constitute a significant obstacle to their use. Of the patients currently being treated, women represent nearly 70%. While early studies have noted sex differences in the response to these drugs, the nature of these differences remain poorly characterized. Using real world electronic medical record (EMR) data, we find that women experience more than double the rates of persistent nausea and vomiting when prescribed GLP1R agonists. To investigate this sex difference in greater detail, we developed novel, species-specific *in vivo* phenomic assays to quantify aversive behaviors. In both mice and rats, aversive responses to either semaglutide or tirzepatide were greater in females than males. To investigate the basis for this difference, we constructed a mouse single cell transcriptomic atlas of body and brain regions most relevant to the action of GLP-1. Using this atlas we find that multiple neuronal cell types involved in the processing of aversive stimuli and nausea had higher GLP1R expression in females than males. Heightened susceptibility of females to the aversive effects of GLP1R agonists could therefore involve increased activation of these brain circuits. Finally, we demonstrate that in mice, both the efficacy and tolerability of GLP1R agonists vary with the phase of the estrous cycle, being highest during proestrus (when estrogen levels peak) and lowest in diestrus (low estrogen levels). Similarly, we report that higher circulating estrogen levels in humans is associated with heightened risk of nausea and vomiting among women taking a GLP1R agonist. Based on these findings, we anticipate that women will continue to be disproportionately impacted by the adverse effects associated with all members of this drug class. Research to better understand and ultimately mitigate this heightened susceptibility is an important priority for new drug development in this area, and novel approaches to model the impact of endogenous hormone signaling will be critical to developing better treatments for more people.

## Introduction

GLP1R agonists have become one of the most impactful drug classes of the last century. Roughly 12%^2^ of the US population have tried a GLP1R agonist and this number is projected to rise substantially over the next decade. Patients taking these drugs routinely achieve >15% weight loss that can be sustained so long as the drug is continued ^3–8^. Yet side effects of these drugs–particularly nausea and vomiting–are common in both clinical trials^9^ and real world data^10^. Approximately 40% of people who have been treated with a GLP1R agonist and stopped cite these side effects as the reason they discontinued taking the medication^11^.

GLP1R agonists mediate their effects on food intake, body weight and metabolic processes^12–15^, by activating GLP1Rs expressed by specific cell types in the brain, pancreas and other peripheral tissues^16–19^. Several studies have highlighted that some of the effects of GLP1R agonists are sex-dependent^20–29^. In the current work, we sought to specifically test whether the ratio of efficacy to tolerability of GLP1R agonists is less favorable for women than men, and to use preclinical animal studies to investigate underlying mechanisms. We were specifically interested in the hypothesis that variation in the circulating estrogen level is a determinant of responsiveness to these drugs. This hypothesis is supported indirectly by evidence that both serum GLP-1 levels and GLP1R expression varies with estrous cycle phase across multiple tissues, although the specific cell types involved have not been characterized ^30–33^. Taken together this would suggest that the total therapeutic profile–effects and side effects–of GLP1R agonists may fluctuate during endogenous hormonal cycles.

In this paper we draw from human electronic medical records, rodent behavioral studies, and single cell RNA sequencing to uncover a role for endogenous sex hormone signaling in mediating the differences in efficacy and tolerability between sexes that are observed with GLP1R agonists. Our findings demonstrate that a broader, systems-level understanding of biology is required to build future treatments for complex metabolic disorders such as obesity.

## Results

### Efficacy-to-tolerability ratio is lower for women on GLP1R agonists

Approximately 70% of people taking a GLP1R agonist for weight loss are female (Fig. 1A) across a broad range of ages (median female = 52.8, male = 56.9). Several clinical trials^22,34–37^ have previously found that women experience both greater weight loss and a higher incidence of adverse events when taking GLP1R agonists in clinical trials. Further work analyzing clinical trial data has demonstrated that these effects remain after taking into account exposure levels (i.e. circulating levels of drug)^38,39^. Beyond this, however, the extent to which inherent, sex-specific biological variables impact responsiveness to this drug class is unknown.

**Figure 1.**
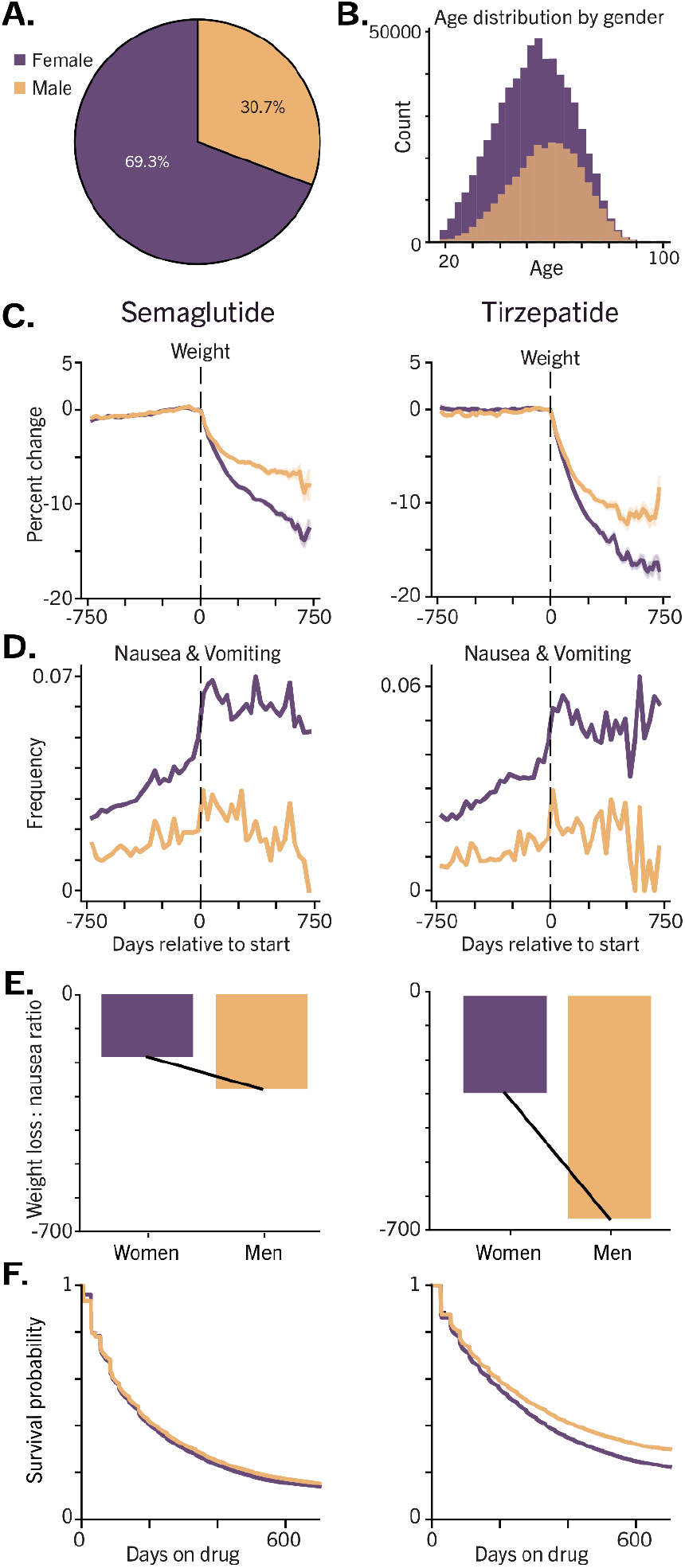
Women respond differently to GLP1R agonists. **A)** 69.3% of patients taking semaglutide or tirzepatide are women. **B)** Histogram of ages for 581,000 patients taking semaglutide or tirzepatide. **C)** Women lose a higher percentage of body weight when taking GLP1R agonists. **D)** Women experience 2.5-fold higher rates of nausea and vomiting on semaglutide and tirzepatide. **E)** Women experience a lower ratio of weight loss to nausea and vomiting than men. **F)** Women are more likely to discontinue both semaglutide and tirzepatide. semaglutide = 1.04 (women are 4% more likely to discontinue at any given point) tirzepatide = 1.16 (women are 16% more likely to discontinue at any given point) p<.005 for all using Cox Proportional Hazard model.

To address this knowledge gap, we used real-world medical record data to investigate sex differences in either efficacy or tolerability outside the clinical setting. We find that for both semaglutide and tirzepatide, women lose more weight (Fig. 1C) and experience a 2.5-fold higher rate of nausea and vomiting than men (Fig. 1D). We also observed that the initial rates of nausea and vomiting for women did not decrease over time for up to two years (Fig. 1D)^40,41^. When comparing the ratio of weight loss to adverse event rates we find that women experience disproportionately more adverse events (i.e. nausea and vomiting) than men per kilogram of weight lost (Fig. 1E). Furthermore this efficacy-to-tolerability ratio appears to be significantly lower for tirzepatide than semaglutide. While correlative data such as these cannot establish causality, the observation that women are also more likely than men to discontinue these medications strengthens the interpretation that the efficacy-to-tolerability ratio for these drugs is lower for women (Fig. 1F).

### Efficacy-to-tolerability ratio is lower for female rats on GLP1R agonists

As a first step to investigate biological mechanisms underlying sex differences in the response to GLP1R agonists, we administered a single dose of semaglutide or tirzepatide in male and female Wistar rats across a wide range of doses (0.0003 mg/kg up to 0.3 mg/kg; IP) and measured food intake over the subsequent 24 hours. As expected, both drugs induced a dose-responsive suppression in food intake that scaled with escalating doses. After normalizing measured food intake to the starting body weight of each animal, we observed no significant sex differences were observed (Fig. 2A). In rats, therefore, the appetite suppressing effects of these drugs do not show a sex bias.

**Figure 2.**
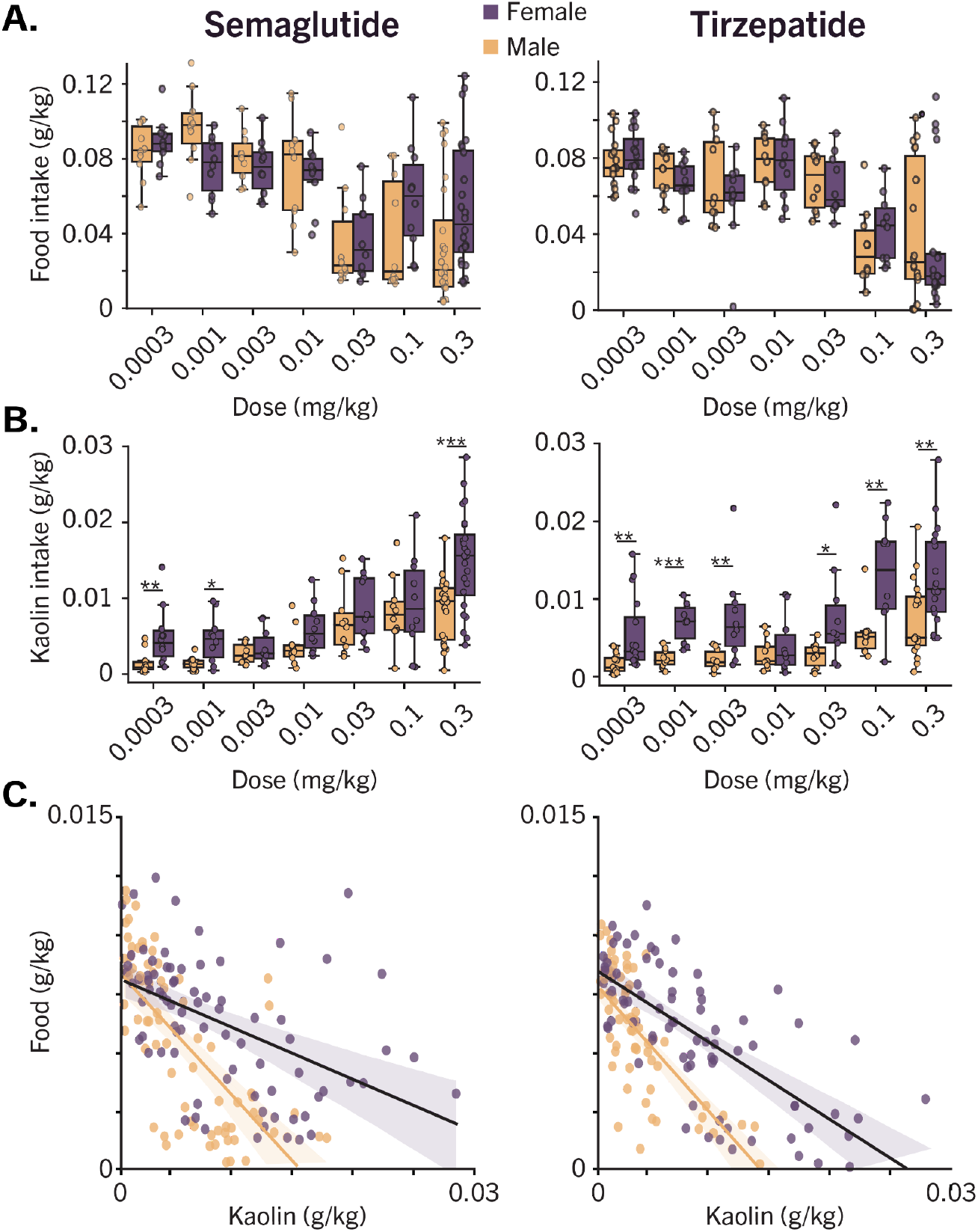
Female rats respond differently to GLP1R agonists. **A)** Food intake suppression is not different between male and female rats. Food intake each day was normalized to animal body weight for semaglutide and tirzepatide. 0 out of 14 dose levels showed a significant difference between males and females. **B)** Pica induction is significantly different between male and female rats. Kaolin intake was normalized to animal body weight for semaglutide and tirzepatide. 9 out of 14 dose levels showed a significant difference between male and female rats. Kruskal-Wallis followed by Mann-Whitney U-test with Bonferroni correction. Semaglutide: H-statistic 13.38, p = 2.5e-4, Tirzepatide: H-statistic 32.97, p = 9.4e^9. *p<.05, **p<.01, ***p<.001. **C)** Slope of food-to-clay intake measures for semaglutide and tirzepatide. The slope of food-to-clay intake changes is shallower for female rats.

Ingestion of non-nutritive clay, also known as ‘pica’, is a commonly used measure of aversion in rodent models^42–47^. In this study, we also provided access to non-nutritive kaolin clay blocks and measured effects of both drugs on clay intake over 24 hours after a 7 day habituation period. As expected, both semaglutide and tirzepatide elicited a dose-dependent increase of clay intake, and for 9 out of the 14 dose/drug conditions, the effect was significantly greater in female than male animals (Fig. 2B). Comparing the ratio of food intake suppression to clay intake across the groups revealed that for all doses for each drug, clay intake was significantly increased in female rats per gram of food intake suppression (Fig. 2C). For a given degree of appetite suppression, therefore, female rats exhibit heightened sensitivity to the aversive properties of these drugs.

### Efficacy-to-tolerability ratio is lower for female mice on GLP1R agonists

Distinguishing between aversive and non-aversive effects is challenging if one relies solely on measures of food intake, body weight, or kaolin clay intake. To address this limitation, we developed a rodent *in vivo* phenomic screening platform (Fig. 3A). This platform starts with a custom “ home cage” where mice can live continuously, but which is outfitted with an array of continuous sensors, a running wheel, and food and water dispensers that are easily monitored. This platform also allows continuous video monitoring (Fig. 3B) in addition to measuring other physiological variables. The information gathered from this approach exceeds that which is captured with a standard metabolic cage but is ∼1/5th the cost, making it easily scalable (Fig. 3C). Using these boxes to monitor C57BL/6 mice following administration of semaglutide, tirzepatide, or vehicle we observed dose-responsive effects not only on “ standard” variables such as reduced locomotion (Fig. 3D), but also on more granular aspects of mouse behavior with each animal generating ∼2.5 million raw datapoints (∼30 FPS) over each 24-hour observation period (Fig. 3E).

**Figure 3.**
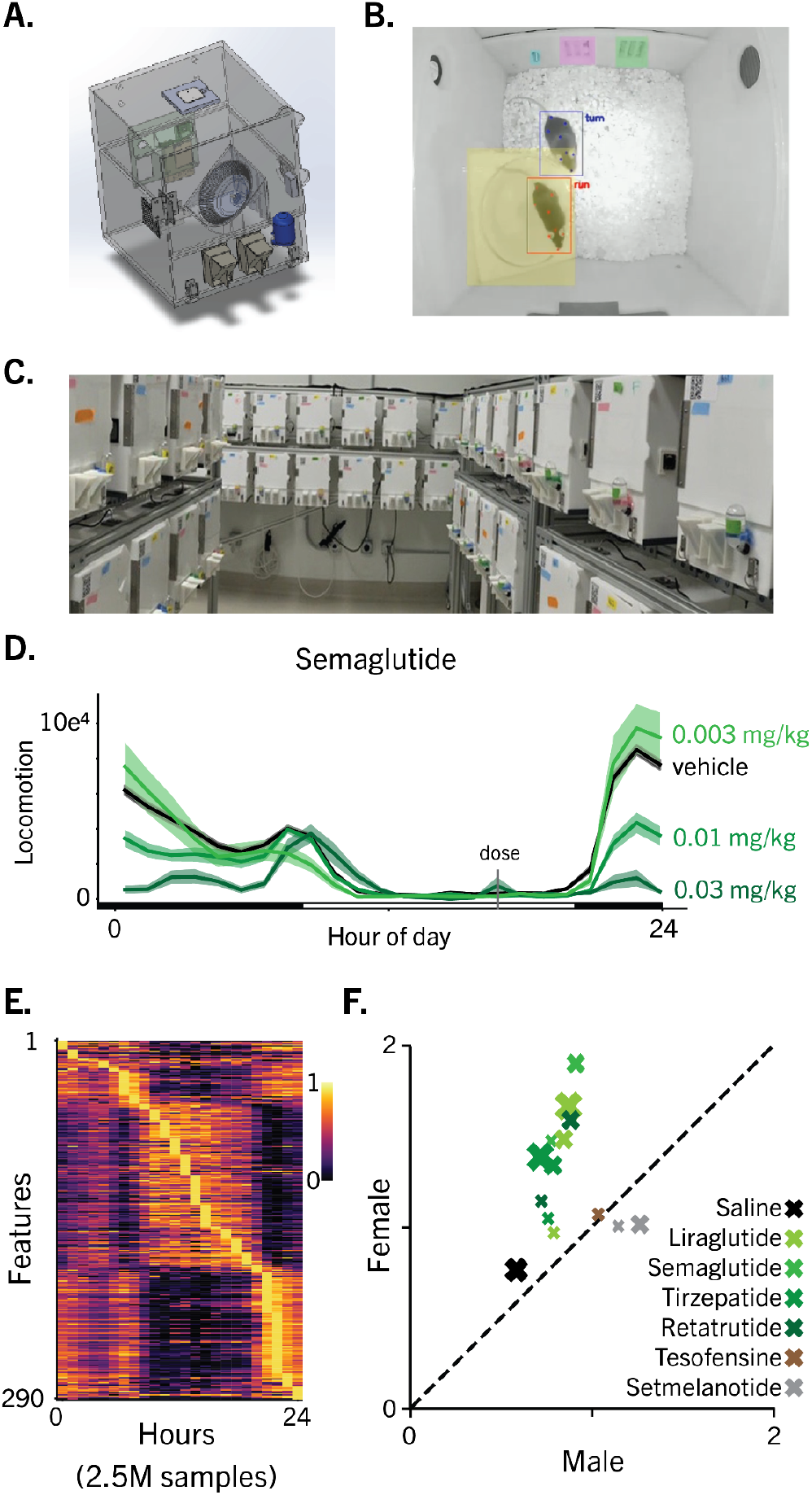
Female mice respond differently to GLP1R agonists. **A)** Box design. Olio has developed a scalable *in vivo* phenomic home cage. Animals have access to two food hoppers, water, a running wheel and are co-housed in pairs. A variety of sensors capture detailed behavioral and physiological patterns from each animal, 24 hours per day. Top down camera view of two mice being tracked. **C)** *In vivo* phenomic homecages are developed for ergonomic and efficient technician access, resulting in workflows 5.8 times faster than standard vivarium solutions. **D)** Example traces of locomotion for cohorts of mice (N>16; 50/50 M/F) administered saline or semaglutide at varying doses. **E)** Olio’s *in vivo* phenomic boxes capture 2.5 million datapoints per mouse per day. **F)** Female mice administered GLP1R agonists display more behavioral and physiological abnormalities than males given the same dose. Icon sizing indicates low, medium, or high dose for each GLP1R agonist. N >= 8F/8M for each drug, N=600 for saline. This sex difference is not observed for non-GLP1R targeting obesity therapeutics such as Tesofensine and Setmelanotide.

Using these boxes in balanced cohorts of female and male C57BL/6 mice revealed a systematic bias wherein drug-treated females deviated from normal (non-injected) sex-matched cohorts than male animals (y-axis is females, x-axis is males). This much more granular behavior analysis tool thus extends our earlier finding of increased sensitivity to the aversive properties of GLP1RAs among female rats based on pica (Fig. 2C). Importantly, there was no sex bias in mice receiving no treatment, saline or a therapeutic dose of the obesity drugs tesofensine or setmelanotide (Fig. 3F). Thus, the behavioral phenotype induced by GLP1R agonist administration is relatively specific in female mice.

### GLP1R expression differences between females and males

Our finding of a consistent sex difference in the adverse effects of GLP1R agonists (after adjusting for differences in weight loss efficacy) across three species raises the possibility of inherent, sex-dependent differences in responsiveness of GLP1R-expressing neurocircuitry. To investigate this hypothesis, we employed an unbiased, discovery-based approach to identify sex differences in GLP1R expression across 48 female and 48 male C57BL/6 age-matched mice. Using scRNAseq, we analyzed brain regions implicated in the actions of these drugs (AP/NTS, PVN, PBN, LS, PG, ARC/DMH/LH), as well as white adipose tissue and liver. In total we collected 860,190 nuclei that passed standard quality metrics (Fig. 4A).

**Figure 4.**
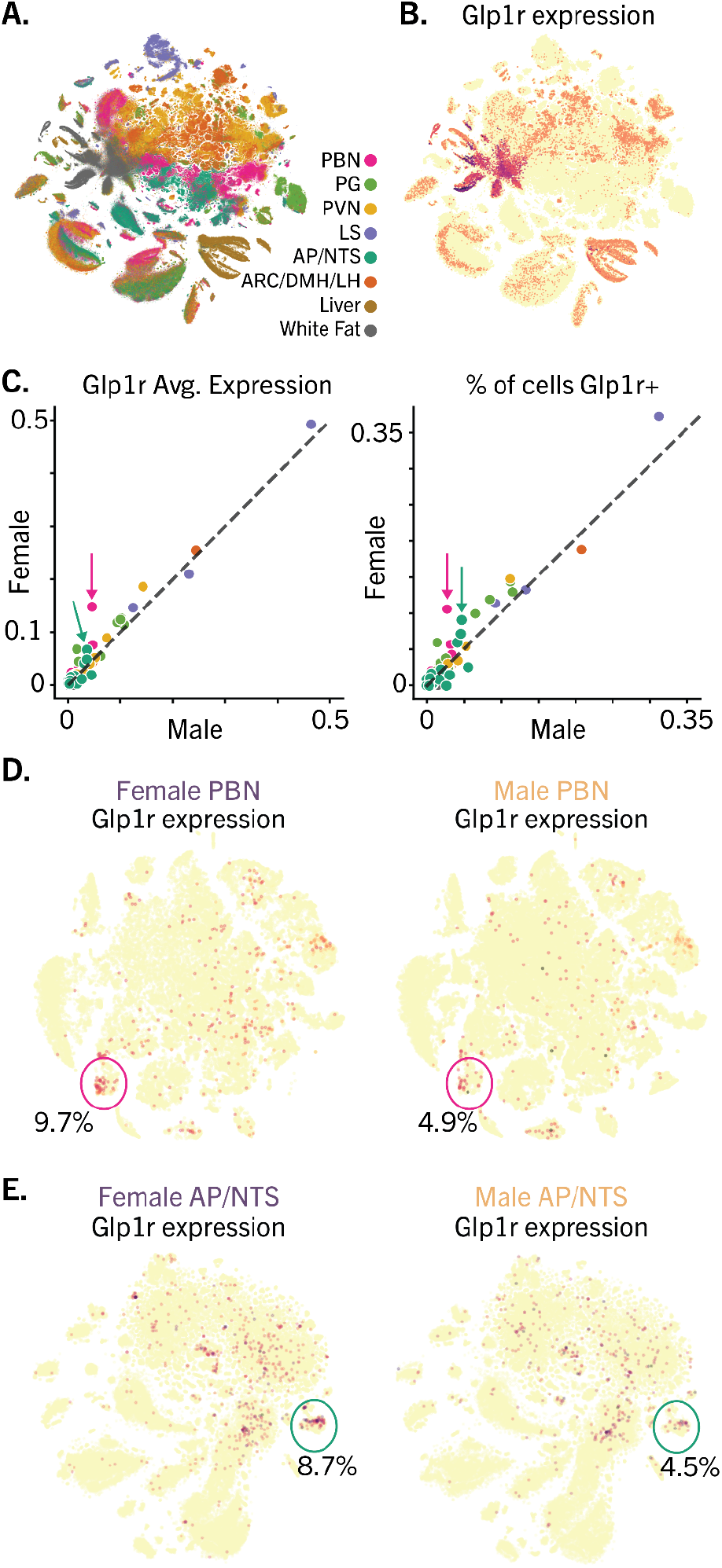
GLP1R is expressed differently in specific cell types in the mouse. **A)** Olio has generated an scRNA-seq atlas of 8 tissues with relevance to GLP-1 signaling. 860,190 passed QC across 8 tissues from a total of 48 female and 48 male C57BL/6 mice. **B)** GLP1R is broadly expressed throughout tissues of the mouse body and brain. **C)** *Left:* For each cluster in our data, we calculated the mean GLP1R expression for cells coming from females (y-axis) and from males (x-axis). *Right:* For each cluster in our data, we calculated the percent of cells with >1 GLP1R transcript for cells coming from females (y-axis) and from males (x-axis). One cluster of parabrachial neurons displayed a >2-fold difference in GLP1R expression between females and males. **D)** Heatmap of Glp1r expression across PBN clusters. One cluster of PBN Glp1r+ cells shows differential expression of Glp1r between sexes. **E)** Heatmap of Glp1r expression across AP/NTS clusters. One cluster of AP/NTS Glp1r+ cells shows differential expression of Glp1r between sexes.

As expected, GLP1R expression was widespread in various cell types throughout the body (Fig. 4B). At the cell cluster level (clustered within tissue type), females tended to express modestly higher levels of *GLP1R* mRNA when compared to the same tissues from males, a difference that in several clusters achieved statistical significance (Fig. 4D). Among brain areas exhibiting these differences are the PBN and AP/NTS, brain areas implicated as mediators of nausea, nociception, visceral malaise and other responses to aversive stimuli (including reduced food intake)^48–51,51–61^. Our finding of increased expression of *GLP1R* mRNA in several cell clusters in the female brain (Fig. 4D,E) offers a feasible and testable hypothesis to explain their heightened sensitivity to the feeding and aversive properties of GLP1R agonists compared to males.

### Aversion induced by GLP1R agonists varies across the reproductive cycle

The inhibitory effects of estrogen on food intake and body weight are well documented ^62,63^, including evidence of an enhanced feeding response to GLP1 ^31,64–66^. We therefore hypothesized that the aversive properties of GLP1R agonists vary with the phase of the estrous cycle. To test this hypothesis, we again used our *in vivo* phenomic screening platform. Specifically, we sought to determine the impact of estrous cycle phase (proestrus, estrus, metestrus and diestrus) on the response to a single dose of semaglutide (0.01 mg/kg) over a 24-h period of observation. A total of 80 female C57Bl6 mice were co-housed in pairs and habituated for at least 10 days prior to study. On the day of semaglutide injection, estrous cycle phase was determined for each mouse (Fig. 5A; see methods). Animals were then sorted post-hoc into 4 groups based on estrous phase, and both body weight and behavioral responses to this dose of semaglutide were compared (Fig. 5C). Strikingly, female mice treated during diestrus (when estrogen levels are low) lost less than half of the amount of weight lost by mice treated during proestrus (when estrogen levels peak). Overall, estrous phase explained 17.8% of the variance in semaglutide-induced weight loss, a highly significant effect size on par with genetic background or disease comorbidity ^67^.

**Figure 5.**
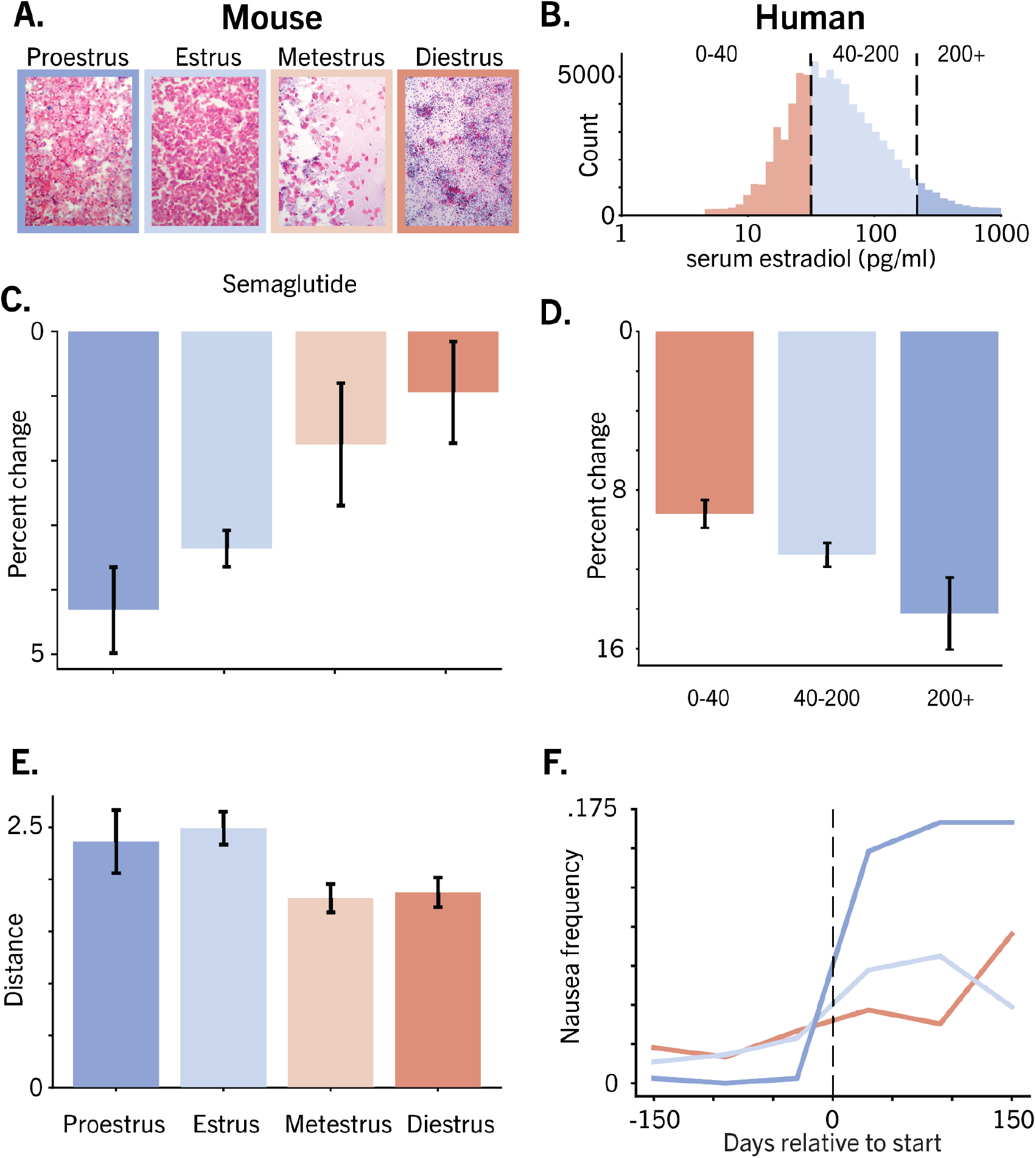
Efficacy and tolerability of GLP1R agonists are modulated by estrous phase and estradiol levels in mice and humans. **A)** Mouse estrous tracking **B)** Histogram of human estrogen levels in EMR data. **C)** Weight loss from GLP1R agonists is dependent on estrous phase in mice. **D)**. Weight loss from GLP1R agonists is dependent on estradiol levels in humans (p=.004, binomial GLM effect between pre/post bins x estrogen level). **E)** GLP1R agonists induce adverse events which are dependent on estrous phase in mice. **F)** GLP1R agonists induce nausea and vomiting which are dependent on estradiol levels in humans. Kruskal-Wallis one-way ANOVA with Dunns post-hoc (H-statistic: 10.84, p=0.004, 1:2 p=.015, 2:3 p=.097, 1:3 p=.002)

However, body weight measurements alone do not allow for interpretation of whether such estrous phase dependent effects were due to changes in appetitive signaling, aversive signaling, or both. To distinguish between these variables we also measured behavioral and physiological parameters (i.e. body temperature, and CO2 levels) from each mouse and evaluated how far these variables diverged from animals who were not dosed with semaglutide. Consistent with the impact of estrous phase on the weight loss response, semaglutide-induced behavioral disruption was also estrous phase-dependent, being greatest during proestrus (Fig. 5E), when estrogen levels peak.

A key question raised by these findings is whether in women, the phase of the menstrual cycle similarly impacts responsiveness to GLP1R agonists, but unfortunately, reliable information regarding menstrual cycle phase could not be deciphered from our EMR data set. Taking an alternative approach, we asked whether elevated serum estradiol levels are associated with enhanced responsiveness to GLP1R agonists in women. We identified 2337 women where two events occurred within 100 days in their medical records, 1) a blood sample had been drawn and serum estradiol measured, and 2) treatment with either semaglutide or tirzepatide was initiated (Fig. 5B). We then separated patients into 3 groups based on measured levels of estradiol (Fig. 5B) and examined the reported body weight change observed over the 100 days following the initiation of either drug. Our finding that the circulating estradiol level was a significant predictor of weight loss offers further evidence that estrogen enhances responsiveness to these drugs (Fig. 5D). Furthermore, when rates of nausea and vomiting were examined in 298 patients that had estrogen measured in the 10 days prior to GLP1R agonist initiation, we found that higher serum estradiol levels were clearly predictive of increased rates of nausea and vomiting in this cohort (Fig 5F). These findings have prompted ongoing studies to disentangle relationships between menstrual cycle phase, serum estrogen levels and the ratio of efficacy to tolerability for GLP1R agonists.

## Discussion

Despite their unparalleled efficacy for both obesity and type 2 diabetes, adherence to GLP1R agonists for obesity or type 2 diabetes is limited by nausea, vomiting and other side effects. Indeed, most patients discontinue the use of these drugs within one year, and nausea and related GI side effects are often cited as the reason.

Here, we report that for a given degree of weight loss, nausea and vomiting are more frequent in women than men taking these drugs. We also show in preclinical studies that GLP1R expression is increased across multiple brain areas implicated in the processing of aversive stimuli. We further demonstrate that in female mice, sensitivity to the aversive properties of these drugs is estrous cycle-dependent, with responsiveness being amplified during proestrus, when circulating estrogen levels peak, and minimized during diestrus, when estrogen levels are low. Finally, we show that in women taking either semaglutide or tirzepatide, elevated circulating estradiol levels are predictive of nausea and vomiting risk. We conclude from these collective observations that across species, females are more sensitive than males to the aversive effects of GLP1R agonists. Accordingly, we anticipate that the many members of this drug class (including various dual- and tri-agonists) currently under development will continue to drive adverse effects disproportionately in women. Given the large number of women for whom these drugs are currently prescribed, these findings are of immediate clinical relevance and they heighten the need to better understand and ultimately mitigate this heightened susceptibility.

A key goal of future work is to identify mechanisms underlying this sex difference. One model supported by our data proposes that in at least some brain areas, the neuronal response to GLP1R activation is amplified by estrogen. Consistent with this model is our finding that in female mice, neuronal GLP1R expression is increased in brain areas known to reduce food intake and promote aversive responses, including the AP, NTS and PBN. It is now well established in murine models that activation of GLP1R-expressing neurons in these brain areas is both necessary and sufficient to explain the feeding and aversive properties of GLP1R agonists ^68,69^. Specifically, projections from GLP1R-expressing neurons in the AP and NTS to hypothalamic nuclei involved in energy homeostasis drive food intake inhibition. In addition, some of these GLP1-responsive neurons also project to the PBN, a brain area well known for detecting and responding to aversive stimuli. Consequently, activation of these neurons reliably reduces intake while also inducing “ sickness behavior” ^53,58^. Furthermore, some PBN neurons themselves express GLP1R, raising the possibility of a more direct mechanism linking the action of GLP1R agonists to nausea and vomiting.

With this background, the question becomes how increased GLP1R signaling in these brain areas might explain sex differences in responsiveness to GLP1R agonists. One possibility is that estrogen effectively raises the circulating level of GLP1R agonists. Another, supported by our data, is that estrogen increases the responsiveness of hindbrain GLP1R-positive neurons to this drug class. A related possibility is that some hindbrain neurons express both the estrogen receptor and GLP1R, and that activation of the former (by estrogen) amplifies signal transduction of the latter (by a GLP1R agonist). Yet another scenario involves alternative circuits. For example, recent work points to an estrogen-sensitive neurocircuit extending from hypothalamus to habenula that shapes aversive states ^70^; whether and how this circuit might be impacted by GLP1R activation is unknown. Each of these potential mechanisms is testable in mice with currently available technologies. A related question of interest is whether the ability of GLP1R agonists to lower rates of diabetes complications (affecting heart, liver, kidney and others) are also sex-dependent.

### Immediate steps to improve therapeutic outcomes with GLP1R agonists

Our results underscore the value of including female subjects in pre-clinical studies, such that sex differences in efficacy or tolerability can be assessed at early stages of the development pipeline. Further, monitoring the estrus phase of female rodents is no longer a prohibitive method for most research institutions. Machine vision algorithms are now available that allow researchers to classify estrous phases based on cytology images^71^ without the need for an in-house expert. By incorporating estrous monitoring into current pre-clinical biomedical research, experimenters can assess differential effects of drugs and doses across estrous phases, which will be critical for understanding how drugs interact with endogenous fluctuations in hormonal state^72^.

Such programs could pave the way for women-specific dosing regimens that could, for example, be altered based on menstrual phase. Widely used menstrual-tracking apps can now make phase-dependent regimens accessible to the general population. We anticipate that by lowering GLP1R agonist dosing during phases associated with the highest estrogen level, the most severe side-effects could be mitigated. In the short-term this could help to mitigate the increase of nausea and vomiting we observed at peak estrogen levels, and thereby reduce discontinuation rates in women, which are higher than in men.

### A new approach to rodent preclinical physiological assessment

Rodents are primarily used in preclinical studies to obtain preliminary results on systemic effects, pharmacokinetics, efficacy, and toxicity before moving on to test drug candidates in larger animals and humans. In many cases, safety and efficacy results from preclinical rodent assays are limited, and often fail to translate to humans^73–75^. Contributing to this limitation, the murine expression of many symptoms are assessed through highly specific behavioral paradigms that are difficult to scale^76,77^. These tasks have been necessary because while certain effects are overt and directly comparable across species, such as weight loss and feeding, others like the subjective experience of pain, are not. To unlock the full translational potential of mouse behavior, we need to move beyond the human-observable.

To this end, we sought to extract as much information as possible from rodent behavioral data and thereby maximize predictive translational power. This involved 24/7 monitoring of pair-housed animals and quantification of behavioral features in their home cages. While the primary efficacy metric – weight loss – was overtly measurable, we took a symptom-agnostic approach to assess safety. Instead of asking how expression of a narrow experimenter-defined set of behaviors is affected by drug exposure, we quantified the extent to which a drug induced abnormal behaviors, whatever that may be. This approach offers unprecedented nuance to our behavioral analysis, allowing the detection of subtle differences across dimensions that might otherwise be missed. A future goal is to increase the translational capacity of our approach by identifying direct, predictive links between behavioral profiles and specific human symptoms and outcomes.

### The future of therapeutics for biologically distinct subpopulations

Our current understanding of the neuronal distribution of GLPRs predicts that GLP1R agonism will always result in nausea and vomiting, unless steps are taken to avert this response. One key question relates to the extent to which neural circuits that drive appetitive and aversive behaviors are dissociable ^68,69^. The consequences of targeting GLP1R across a broad array of cell types highlights a more general principle, that relatively few proteins are expressed in a manner sufficiently selective for optimal drug development. Thus, alternative targeting approaches that are tailored to inherent biological differences should lead to better therapeutics. Drug development programs that take inherent biological diversity into account at the earliest preclinical stage will be able to design better medicines for a larger number of patients.

## Methods

### Animals

Protocols were approved by an external Institutional Animal Care and Use Committee. Male and female C57BL/6 mice were purchased from Jackson Laboratories at age 8 to 12 weeks. Mice were habituated in custom built behavior boxes equipped with a running wheel in a 12/12 light cycle for 10 days and fed ad libitum water, high fat diet (HFD, Research Diets D12492i) and regular chow (Labdiet Picolab 5053) in separate hoppers. Each box housed 2 mice of the same sex, with equal distribution between sexes. After 10 days in the boxes, mice were eligible for drug injection.

### Rat food and kaolin clay studies with GLP1R agonists

Wistar rats aged 6-8 weeks were habituated to housing for one week. Regular chow and kaolin clay were provided during this habituation period. There were 24 rats per drug (48 total), 12 male and 12 female. Rats were assigned to 6 different groups that received a different dose of the same drug on a given day such that all rats received 3 different doses of a given drug. Rats were dosed via IP injection every 3 days. Semaglutide doses were: 0.1, 0.3, 1, 3, 10, 30 nmol/kg, tirzepatide doses were 0.3, 1, 3, 10, 30, 100 nmol/kg. Food and kaolin intake was measured 24 hours after dosing.

### Experiment design

Mice were randomly assigned to an injection group and would receive an IP injection of a different, randomized drug or drug combination with a minimum of 3 days between injections. Post injection, behavior and physiological parameters were monitored via video, thermal camera, CO2 sensor and humidity sensor for 12 hours. Body weights, food weight in the hoppers, and water weight in water bottles were measured just prior to injection. At the end of 24 hours, all weights were measured again. Mice were always injected at the same time of day and total injection volumes were kept below 0.3 ml, with 0.1 ml median injection volume.

### Semaglutide dosing and estrous tracking

Female mice were given an intraperitoneal injection of either semaglutide (n=80) at 0.01mg/kg or saline control (n=80). Immediately prior to injection, vaginal cytological samples were collected and transferred to gel coated glass microscope slides for staining and inspection. Cytological sample collection was performed either using vaginal lavage with sterile distilled water or cotton swabs with sterile saline solution. Samples were stained using Hematoxylin and Eosin then labeled by at least two experts using well-established criteria^78,79^.

In brief, four stages were classified according to the following cytological characteristics. Proestrus is defined by large proportions of small nucleated epithelial cells (SNEs), but low numbers of leukocytes (LEUs), large nucleated epithelial cells (LNEs), and non-nucleated keratinized epithelial cells (AKEs). Estrus is defined by an overwhelming proportion of AKEs, and smaller numbers of SNEs, if any. During metestrus, SNE and LNE numbers increase, AKE numbers decrease, and LEU numbers increase but cluster around SNE and LNEs. Diestrus can be distinguished by the high proportion of LEUs, either clumped or dispersed, with relatively small numbers of other cell types present, if any.

### Behavioral quantification

Video footage of mice was captured using USB cameras positioned on the ceilings of each experimental apparatus. To extract relevant frame-by-frame information from the video data, we employed deep learning models that inferred bounding boxes surrounding each mouse and keypoint locations across various body parts (YOLOv8). These model-derived predictions were then processed to analyze numerous dimensions of mouse behavior. Specifically, we computed features encompassing movement dynamics (e.g., locomotion and turn biases), social interaction metrics (e.g., inter-mouse distances and relative orientations), and action-specific dynamics (e.g., transition probabilities between actions). In total, 291 distinct behavioral features were extracted for each video frame within each experiment.

To generate a summary representation of behavior, behavioral features were then binned into hourly intervals, and the median feature values were computed across the two mice sharing each experimental apparatus. For interpretable comparisons of behavior under drug conditions versus normal conditions, the dimensionality of the behavioral data was reduced from 291 features to 39 using a combination of feature selection and spectral embedding of action-transition matrices.

Behavioral deviation from normal conditions was quantified by assessing the aggregate differences between features in the drug versus no-drug conditions. The distributions of features from normal and drug data were compared using population stability index (PSI) analysis, a symmetric measure of distribution drift whose scores are comparable across different datasets. Next, the median of these scores across features was computed to generate a single deviation value. This deviation value was used as the safety metric, where lower values correspond to safer outcomes.

### scRNA experiment

48 male and 48 female C57BL/6 mice were obtained from Charles River Laboratories at 6 weeks of age. After 3 weeks of acclimation to our animal facility they were grouped into sex matched CTL and HFD groups and fed Research Diets HFD diet (60 kcal% fat, product # D124k) or control diet (10 kcal % fat, product # DS1249) respectively for 12 weeks. At the end of this period, animals were weighed, and their brains were extracted and flash frozen in isopentane, and then immediately sectioned. Punches of the following brain areas were extracted: lateral septum (LS), arcuate nucleus, dorsomedial and dorsolateral nucleus of the hypothalamus (HYP), periventricular nucleus of the hypothalamus (PVN), pontine gray (PG), parabrachial nucleus (PBN), and the nucleus of the solitary tract and area postrema (NTS/AP). Samples from 3 animals were pooled into one tube. Further, we collected the liver and white fat tissue from each mouse. The body tissues were not pooled due to their larger size. Brain and body tissue samples were transferred to Seqmatic, a local single cell scRNA company based in the Bay Area, for nuclei extraction, fixation, barcoding and library preparation using a Parse Megakit. A 300 cycle NovaSeq X25 B sequencing run was performed on the samples. We obtained readouts of 860190 cells from 8 tissues.

### scRNA analysis

Internal scRNA-seq data was processed using standard Parse Bio pipeline. After filtering out low quality cells based on total UMI counts (<500 UMIs), percent mitochondria reads (>=5%) and potential doublets expressing both female- and male-specific marker genes, initial global clustering and UMAP was performed to first separate neurons vs. non-neurons using canonical marker genes (e.g., Slc17a6, Slc32a1, Snap25, Syp etc.) for neurons, Mag for oligodendrocytes, Ntsr2 for astrocytes). Then cells expressing marker genes for two broad cell types (e.g., expressing both neuronal and astrocyte markers) were removed. Clustering, UMAP and tSNE was also performed for each tissue separately to increase clustering resolution.

### Human EMR data

The population of semaglutide and tirzepatide users was selected according to Rodriguez et al., 2024 and a data snapshot created 1/27/2025. First dispenses to a deduplicated patient of semaglutide or tirzepatide in the MedicationDispense table was used to establish medication start date. All doses of both semaglutide and tirzepatide were pooled. Nausea and vomiting events were classified using Observation and Condition tables from the Truveta dataset using SNOMET CT, ICD10M, ICD9M, and LOINC codes: **Nausea** (*SNOMED CT:* 2919008, 16932000, 73335002, 162057007, 386368005, 419219000, 422587007, 698861005). **Vomiting** (*SNOMED CT:* 422400008, 16932000, 1488000, 405166007, 2919008, 698861005, 249519007, 146291000119108, 23971007, 15387003, 18773000, 71419002, 196746003, 8579004, 139337005, 63722008, 246452003, 90325002, 236083006, 49206006, 74621002, 419219000, 294001000119105, 332982000, 73335002, 12860001000004105, 37224001, 162063003, 158420004, 158422007, 275297005; *ICD10CM:* R11.2, R11.10, R11.11, R11.12, R11.1; *ICD9CM:* 787.01, 787.03, 536.2; *LOINC:* 81224-8). Nausea and vomiting events were pooled. Estrogen measurements were obtained from lab result records using CPT and LOINC codes: Estrogen (*CPT:* 82671, 82672, 1011401, 82679; *LOINC:* 2254-1, 53765-4, 53766-2, 2255-8, 34295-6, 2243-4, 2258-2).

Discontinuation of medication was established by determining how much medication a patient had on hand on a given day calculated by the amount of medication for a given dispense event and how much was left at the date of the dispense event from previous dispenses. 60 days without medication was considered discontinuation, and the last day of available medication was considered the discontinuation event.

To determine nausea and vomiting rates after medication start, estrogen measurements taken 10 days prior to medication start were used to bin patients based on estrogen level (Levels: <40 pg/ml, 40-200 pg/ml, >200 pg/ml, n=298 patients). Nausea and vomiting events were binned per 60 days. If a patient discontinued medication in a bin, they would not be included in the next bin. Applying a binomial GLM indicated a significant effect of estrogen level on nausea and vomiting rates (p=.002 for pre/post GLP1R agonist, and p=.004 for pre/post x estrogen range)

To determine percent weight loss, weights were first cleaned according to Rodriguez et al., 2024. Briefly, outlier measurements greater than 30% of the median weight for a given person were excluded. Baseline weight was established by taking the median of all weights in the 60 days prior to medication start. Percent change was calculated by subtracting the baseline weight for a person from a given measurement and dividing by the baseline weight. People were divided into estrogen levels according to the median estrogen level in the 100 days prior to medication start, divided into the same bins as in the previous section (n=2337). We expanded the window to capture how chronic estrogen levels affect weight which is affected over a much longer time period. To calculate weight change at 6 months, percent weight change from 150 to 210 days was binned and averaged. A Kruskal-Wallis one-way ANOVA with Dunns post-hoc demonstrated significant differences between (Kruskal-Wallis H-statistic: 10.84, p=0.004, Dunns test groups 1:2 p=0.015, 2:3 p=0.097, 1:3 p=0.002).

